# Reward Value Is More Important Than Physical Saliency During Bumblebee Visual Search For Multiple Rewarding Targets

**DOI:** 10.1101/2020.10.01.322172

**Authors:** Vivek Nityananda, Lars Chittka

## Abstract

Several animals, including bees, use visual search to distinguish targets of interest and ignore distractors. While bee flower choice is well studied, we know relatively little about how they choose between multiple rewarding flowers in complex floral environments. Two important factors that could influence bee visual search for multiple flowers are the physical saliency (colour contrast against the background) of flowers and the reward value associated with them. We here investigated how these two different factors contribute to bee visual search. We trained bees to independently recognize two rewarding colours that in different experiments differed in either physical saliency, reward value or both. We then measured their choices and attention to these colours in the presence of distractors in a test without reinforcement. We found that bees preferred more salient or higher rewarding flowers and ignored distractors. When the high-reward flowers were less salient than the low-reward flowers, bees were nonetheless equally likely to choose high-reward flowers. Bees were more also more likely to attend to these high-reward flowers, with higher inspection times around them and faster search times when choosing them. When flowers differed in reward, we also found an effect of the training order with low-reward targets being more likely to be chosen if they had been encountered during the more immediate training session prior to the test. Our results parallel recent findings from humans demonstrating that reward value can attract attention even when targets are less salient and irrelevant to the current task.

## Introduction

Animal foraging behaviour is very well studied, but research in this area has not often considered more psychological aspects of foraging such as attention and visual search. Adapting human visual search experiments to investigate visual search in other animals, including bees, has led to an increased understanding of their foraging behaviour and holds promise to become a productive field of research (Dukas and Kamil 2001; Bond and Kamil 2002; Spaethe et al. 2006; Morawetz and Spaethe 2012; Nityananda and Pattrick 2013; Ben-Tov et al. 2015; Orlowski et al. 2015, 2018; Saban et al. 2017). Visual search experiments typically present individuals one target in middle of distractors. Studies have also looked at how attention is deployed when more than one instance of a target type is present (Horowitz and Wolfe 2001) or how attention is divided across multiple tasks (Miller 1982). Fewer studies have looked at visual search for multiple object types or categories that are presented simultaneously (Duncan 1980; Huang et al. 2007; Kristjánsson et al. 2014; Berggren and Eimer 2020). Yet in real life we might well be searching for multiple items at a time, such as say, tomatoes and onions in the supermarket.

In bees, studies of visual search have also focussed on how they choose single targets over others, and we know less about how they search in complex floral environments. In particular, research has focussed on flower constancy, the tendency of bees to specialize on one flower type (Heinrich 1979; Wells and Wells 1983; Waser 1986; Hill et al. 1997). A prominent explanation of flower constancy is that it is a result of a cognitive limitation (Waser 1986; Raine and Chittka 2007), suggesting that bees cannot simultaneously choose multiple flower types. This view has been challenged by recent work showing that if bees are given the opportunity to learn multiple rewarding flower types, they readily approach both types, flexibly switching between the two (Nityananda and Pattrick 2013). In fact, they appear to be able to learn several different target types and this ability is reflected by changes in the neural structures in their brains (Li et al. 2017). Bees can thus clearly select multiple flowers simultaneously, but we still do not know the factors influencing their choices between these flowers.

In humans, several factors are known to influence visual search (Wolfe and Horowitz 2004; Wolfe 2020), but two broad processes have typically been identified as fundamental. These are often classified as bottom-up and top-down visual search (Johnson and Proctor 2004). Bottom-up processes involve an involuntary, rapid capture of visual attention by physically salient stimuli. Top-down processes are more deliberate and guided by the goals of an immediate task. More recently, a third category of processes has been proposed involving the influence of search history (Anderson et al. 2011a; Awh et al. 2012; Anderson 2019; Theeuwes 2019). The most prominent examples of these processes have focussed on the role of reward value (Anderson et al. 2011a, b). Target stimuli that are relevant or monetarily rewarding in one task have been shown to capture visual attention even when they are irrelevant to a subsequent task and not physically salient (Anderson et al. 2011b; Bourgeois et al. 2017; Bucker and Theeuwes 2017). The capture of visual attention in these cases is also involuntary and rapid, as is typically seen in response to physically salient stimuli. Thus, visual search and attention can be influenced by three different processes dependent on physical saliency, current goals and search history.

The physical saliency of flowers as measured by their colour contrast against the background influences flower choice in bees (Lunau 1990; Lunau et al. 1996; Goulson 2000) and would also be expected to influence visual search and attention. Goal-driven visual search is more difficult to study in bees given the impossibility of providing verbal instructions to set goals for them. One way of specifying targets for the bees is to reward certain targets compared to others and reward value (sucrose concentration) does influence flower choice in bees (Benard et al. 2006; Avarguès-Weber and Giurfa 2014). However, this better resembles reward-based visual search than goal-directed search. Both reward and physical saliency could therefore influence visual search in bees. A bee might, however, simultaneously encounter flowers with differing saliency and reward and it is not yet known how these different factors could interact and influence visual search. In this study, we therefore ran a series of experiments to test how saliency and reward influence bee visual search for two simultaneously rewarding target types.

## Methods

### Bees

We obtained the bees from a commercial supplier (Syngenta Bioline, Weert, The Netherlands) and tagged them with Opalith number tags (Christian Graze KG, Weinstadt-Endersbach, Germany) to allow for individual identification. The bee colonies were transferred under red light to one chamber of a two-chambered wooden nest box (28×16×11 cm length × width × height). The floor of the other chamber was covered with cat litter to give bees an area to discard refuse. The nest box was connected through a 24.5 cm long transparent Perspex tunnel to an arena consisting of a wooden box (100×60×40 cm length × width × height) covered with a UV-transparent Plexiglas lid. The bees could enter this arena to forage for sucrose solution. The floor of the arena was covered with green card and the illumination was provided from above using two twin lamps (TMS 24 F with HF-B 236 TLD (4.3 kHz) ballasts; Philips, The Netherlands) fitted with Activa daylight fluorescent tubes (Sylvania, New Haven, UK). Pollen was provided directly into the colony on alternate evenings.

### Spectral Measurements

We measured the reflectance spectra of the artificial flowers using an Avantes AvaSpec 2048 spectrophotometer (Anglia Instruments Limited, Soham, UK) with a deuterium-halogen light source, relative to a BaSO_4_ white standard. To account for the difference between spectral sensitivity in humans and bees, we converted the spectra of the targets into a bee-specific hexagonal colour space (Chittka 1992) incorporating the spectral sensitivity of bumblebee photoreceptors (Skorupski et al. 2007), the spectral reflectance of the background and the spectral distribution of the lights used. The colour hexagon has three vertices corresponding to maximal excitation of each of the bee photoreceptors, which are tuned to green, blue and ultraviolet (UV) light (Chittka 1992). Three further vertices correspond to colour mixtures resulting from approximately equal excitation of two spectral receptors. The Euclidean distance between the centre of the hexagon and each of these vertices is 1 and colour distances greater than 0.1 are well distinguished by bees without special training procedures (Dyer and Chittka 2004a). Once plotted in this colour space (Fig 1), the colour loci can be used to calculate the distances in colour space between pairs of colours, thus indicating the perceptual discriminability of the colours. All measures of colour differences between the artificial flowers used in our experiments are provided in Table S1.

**Fig. 1.**
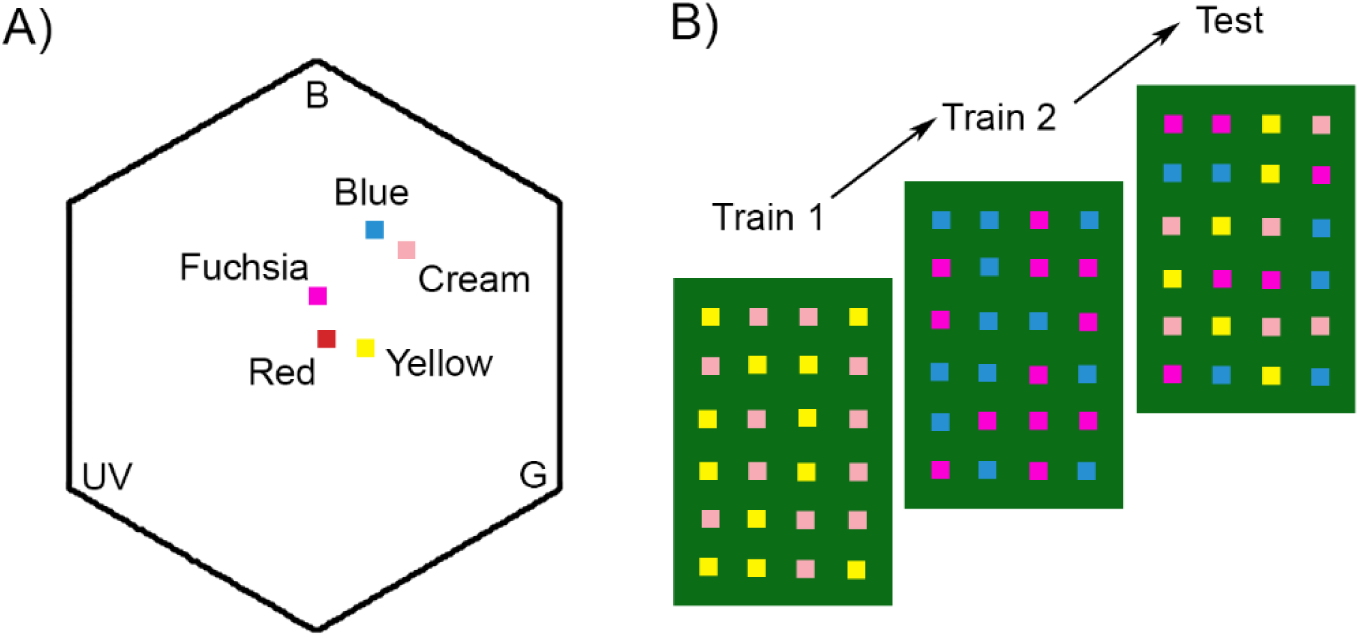
A) Colour loci of the artificial flowers used across all experiments in the colour hexagon. Three of the vertices correspond to maximum excitation of the bumblebee photoreceptors sensitive to ultraviolet (UV), blue (B) and green (G). The angular distance from the centre represents the hue as perceived by the bee. Distances between points indicate the hue discriminability. The distance between the centre and any vertex is 1 and colours that differ by distances above 0.1 are easily distinguishable. B) Example training and test protocol used in the experiments. Bees were trained on one rewarding and one non-rewarding colour in each training session (Train 1 and Train 2) and tested without reinforcement with all four colours in the test session (Test).

### Pretraining

We trained colour-naïve foraging bees to forage from square transparent Perspex chips (side: 25 mm, thickness: 5 mm) that served as artificial flowers (henceforth “flowers”). Each flower had a well in the centre into which rewarding (sucrose solution) or non-rewarding (water) liquids could be placed. After bees learned to approach these flowers, we placed them on glass vials (4 cm tall, 1.5 cm in diameter) and trained the bees to feed from them when they were arranged in a 6 × 4 horizontal grid, with vials placed 15 cm apart. In this grid, twelve randomly chosen flowers had 12 μl of 50% sucrose on them and the others remained empty. The positions of rewarding and non-rewarding flowers in all experiments were determined using the random number generator function RAND() in Microsoft Excel®. Once the bees had foraged on this grid for three bouts, we commenced training.

### Training

In each experiment we trained bees from three different colonies on two visual discrimination tasks. The tasks involved bees having to discriminate target flowers of one colour from distractor flowers of another colour. The flowers consisted of coloured Perspex chips placed in a grid as described above. The 12 target and 12 distractor flowers were placed in positions on the grid that were randomly chosen for each bout of the training. Target flowers held 12 μl of sucrose solution, while distractor flowers held 12 μl of distilled water. Flowers were not refilled during a given training bout. Each bee was individually trained on one of these tasks until it reached a success criterion of 80% correct choices out of the last 20 choices made. Choices were recorded when the bee probed the flowers for reward and bees could revisit flowers in all experiments. Between training bouts, we cleaned the flowers with 99% ethanol to remove scent markings, and subsequently with water to remove any traces of ethanol.

Once a bee successfully completed one training task, it was presented with another task consisting of target and distractor flowers with different colours from those in the first task. The order in which each of these tasks were presented was alternated between bees. The exact details of the colours and reward used are given below. Bees thus sequentially learnt two different rewarding colours.

### Experiment 1: How does physical saliency influence bee visual search?

In this experiment, twenty bees were trained on one physically salient target and one less salient target in separate discrimination tasks. For ten of these bees, one of the two tasks had Blue as the rewarding colour and Cream as the non-rewarding colour. The other task had Fuchsia as the rewarding colour and Red as the non-rewarding colour. Both target colours provided a reward of 50% sucrose solution (v/v). The experiment was replicated with another ten bees using a different set of colours. In this replication, the rewarding colours were Fuchsia and Red, while the distractors were Cream and Yellow respectively. This replication ensured that Fuchsia, the less salient colour (compared to Blue) in the first combination was the more salient colour (compared to Red) in the second combination of colours (Fig 1).

### Experiment 2: How does reward value influence bee visual search?

In this experiment, 15 bees were trained on one high-reward target and one low-reward target in separate discrimination tasks. One of these tasks had Blue as the rewarding colour and Fuchsia as the non-rewarding colour. The other had Cream as the rewarding colour and Yellow as the non-rewarding colour. These colours were chosen as the Blue and Cream colours were close in physical saliency, defined as colour contrast with the background (Table S1). In the two discriminations tasks, one of the target colours had a reward of 50% Sucrose solution (v/v) while the other had a reward of 30% Sucrose solution (v/v). With one exception, the association of high and low rewards with each of the target colours (Blue and Cream) was counterbalanced across all trials as was the order in which bees experienced high and low reward in their two training tasks.

### Experiment 3: How does bee visual search combine reward value and physical saliency?

In this experiment 16 bees were trained on two discrimination tasks. One of these had a high-reward target offering a reward of 50% Sucrose solution (v/v). This target was Yellow in colour and had low colour contrast (i.e. physical saliency) against the background. The distractor in this task was Cream in colour. In the other task, the target offered a lower reward of 30% Sucrose solution (v/v). The target was Blue in colour and had a high colour contrast against the background and the distractor was Fuchsia in colour. The order in which bees encountered each of these tasks was counterbalanced.

### Test

In all experiments, once training was completed, we tested bees on their visual search when faced with multiple targets. We presented the bees with six flowers each of the two rewarding colours they were trained on and six flowers of each of the distractor colours. All flowers in the test were non-rewarding containing 12 μl of distilled water. This prevented reinforcement learning during the test. We noted the choices made by the bees and the order they were made in. The foraging bout of each bee during the test was recorded using a Sony DCR-SR58E Handycam to enable later analysis of the times between the choices. We ran the tests until five minutes were over, or the bee returned to the colony after making at least 12 choices, whichever occurred sooner.

### Data Analysis

For all experiments, we split the choices made by the bees into the different transitions between colours and noted which were switches to different colours and which were constant transitions. We examined the number of constant transitions made before each switch to measure how often bees had runs of constant choices. We then calculated a sequence index for each bee by dividing the number of constant transitions by the total number of transitions (Heinrich 1979). This index is the probability of constant transitions compared to switches. An index close to 1 would indicate that the bees were flowers constant while a value close to 0.5 would indicate that bees switched flowers with every new choice. We used a Wilcoxon rank sum test (α=0.05) to compare the observed number of constant choices with the index values of 1 and 0.5. We also examined how quickly bees made these different choices by comparing the median times taken to make constant choices and switches using Wilcoxon rank sum tests (α=0.05). Since the bees occasionally flew around the arena for extended periods of time without making a choice, we ran an outlier analysis for the times within each category, and excluded data points that were greater or less than 1.5 times the interquartile range prior to the second analysis. Timing data is missing for one bee in Experiment 1 and five bees in Experiment 2 because of the lack of video recordings.

To examine how different factors influenced the proportions of choices made by the bees we ran generalized linear models with the proportion of choices as a dependent variable and the different factors as independent variables. For experiment 1, the independent variables were physical saliency (high or low) and second variable representing the training order. This second variable was a binary variable representing whether the bee first encountered the high saliency target or the low saliency target during the training on visual discrimination tasks. For experiment 2, the independent variables were reward value (high or low) and a second binary variable representing whether the bee first encountered the high reward target or the low reward target during training. For experiment 3, we also had the two independent variables as in experiment 2. In all the models, bee identity was modelled as a random variable and the proportion of choices were modelled as a binomial distribution with a logit link function. We ran models looking for main effects of the independent variables and interaction effects between the variables as well. In this and all other analyses, models were compared using the Akaike Information Criterion (AIC) and the model with the lowest criterion was chosen. The significance of each variable was compared against an α of 0.05.

In experiment 3, we were also interested to see if higher reward could influence bee attention to a target with low physical saliency. We used the positions of the bee during visual search as a proxy for attention. Using the open-source program Tracker (V5.15, ©2020 Douglas Brown, physlets.org/tracker), we perspective corrected each video and tracked the position of the bee in each frame of the video recording during the test phase. We used this to analyse bee behaviour during the first two minutes of the videos. Frames in which it was not possible to spot the bee – either because it flew to the corner of the arena or due to reflections of the lighting-were labelled as missing data. Using the tracked positions of the bees we obtained a map of search behaviour for each bee. We specified zones on these maps corresponding to flower areas and non-flower areas. Flower areas were areas within 2 cm of the flowers. All other areas were non-flower areas. We measured inspection time as the time each bee spent in each of these areas by summing the number of video frames in which bees were present in them and multiplying this by the frame rate (25 frames per second). We compared the inspection time for the different types of targets and distractors. We used a generalized linear model to model this as a binomial variable with a logit link function. As in the analysis above we used reward value, physical saliency and search history as independent variables and bee identity as a random factor. We ran models looking for main effects of the independent variables and for interaction effects between the variables as well.

All statistical analyses were run in RStudio (version 1.2.5033)

## Results

### Experiment 1: How does physical saliency influence bee visual search?

The average time taken for the first and second training bouts on this experiment was 2080.7 (± 1418) seconds and 971.9 (± 366.4) seconds respectively.

Combining results from both flower sets we found that the average proportion of salient target flowers chosen during tests was 0.58 (± 0.13 S.D.) and the average proportion of equally rewarding non-salient targets chosen was 0.37 (± 0.11 S.D.). The average proportion of distractors chosen was 0.06 (± 0.08 S.D.). If bees chose equally between the two targets without choosing any distractors, we would expect an equal proportion (0.5) of both salient and non-salient targets to be chosen. Saliency had a significant effect on the proportion of targets chosen; the proportion of high-saliency targets chosen was significantly greater than the proportion of non-salient targets chosen (GLMM, Effect size estimate: - 0.84, P = 4.3 * 10^−9^, Fig 2A) and the proportion of distractors chosen (GLMM, Effect size estimate = - 3.24, P < 2 * 10^−16^). The low number of choices made to distractors demonstrates that the bees had memorised both types of previously rewarding targets in the training bouts and could recall them in the presence of distractors. The best model that described the data did not include the effect of training order indicating that this was not an important determinant of the proportion of salient targets chosen.

**Fig. 2:**
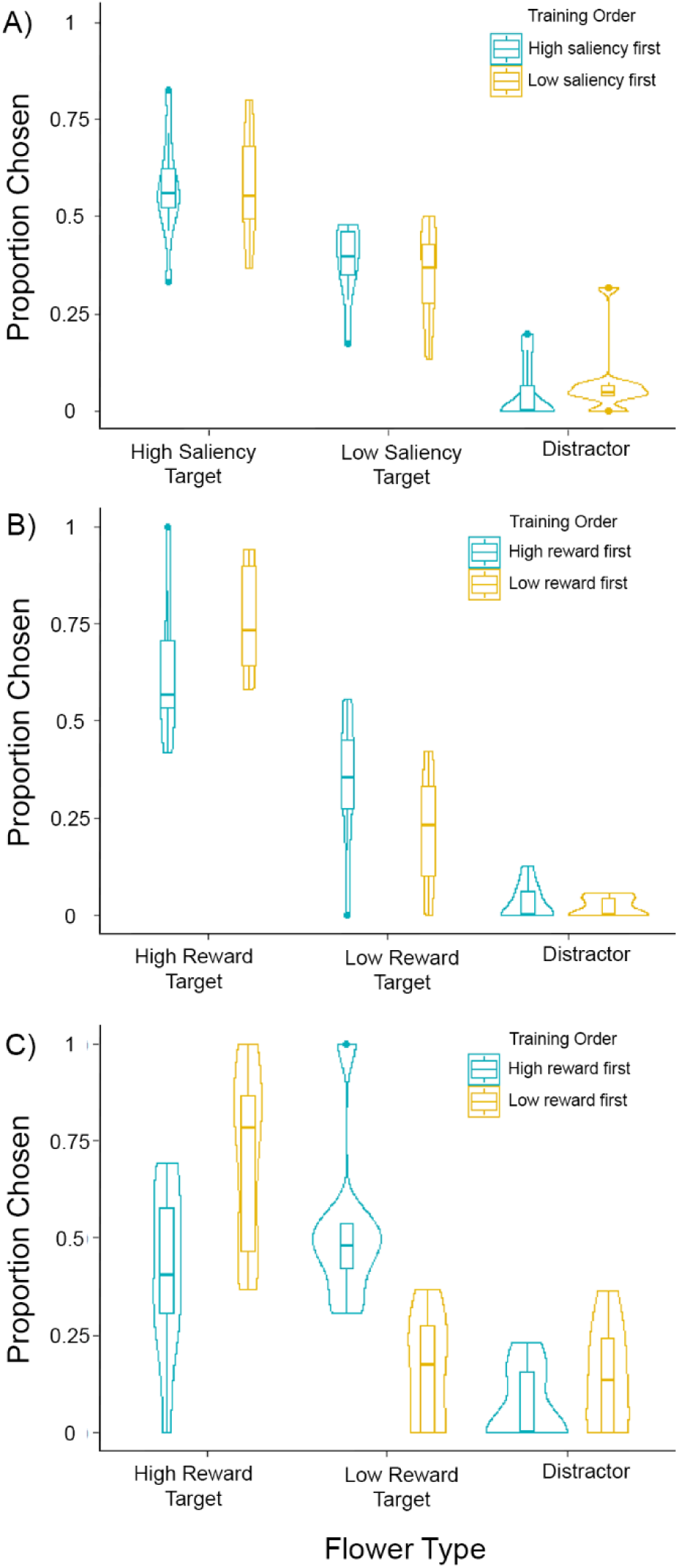
Proportions of different flower types chosen in A) Experiment 1, B) Experiment 2, and C) Experiment 3. Box plots depict the median and the first and third quartiles, the whiskers depict the largest and smallest values that are within 1.5 times the interquartile range from the edge of the boxes. Violin plots overlaid on top of the box plot depict the mirrored density plots of the data.

The average sequence index of the bees was 0.51 (± 0.17 S.D.). An index close to 0.5 indicates equal numbers of constant choices and switches, while an index close to 1 indicates complete flower constancy with no switches. This index was not significantly different from 0.5 (Wilcoxon rank sum test, W = 200, P = 0.1), showing that the bees were equally likely to make constant choices and switches (Fig 3). The times taken for choices between like flowers and transitions between flower types were not significantly different (Wilcoxon rank sum test, W = 13036, P = 0.14). The mean time taken for constant choices was 7.53 (± 4.93 S.D.) seconds compared to a mean of 9.03 (± 7.05 S.D.) seconds for switches (Fig 4A).

**Fig. 3.**
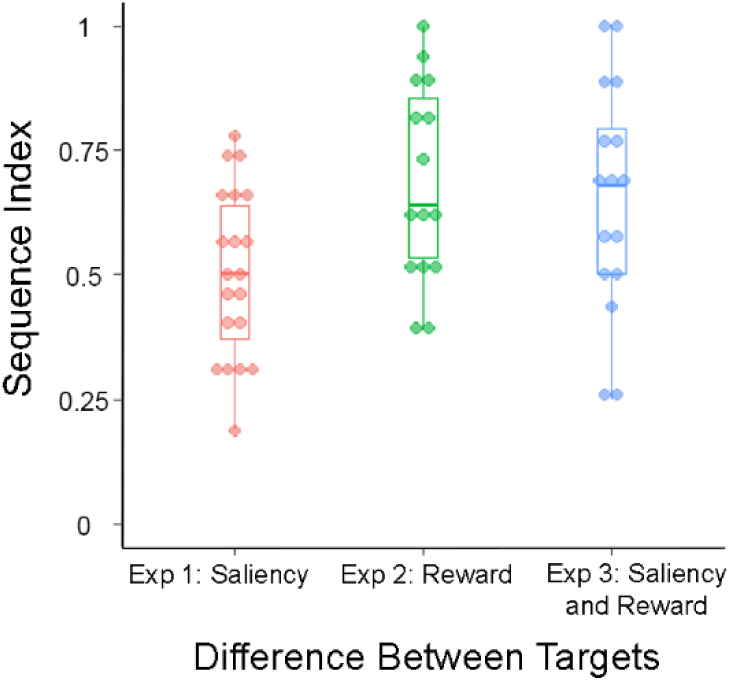
Sequence indices in each of the experiments. Box plots as described for Figure 2. The actual data points are overlaid on top of the box plot.

**Fig. 4:**
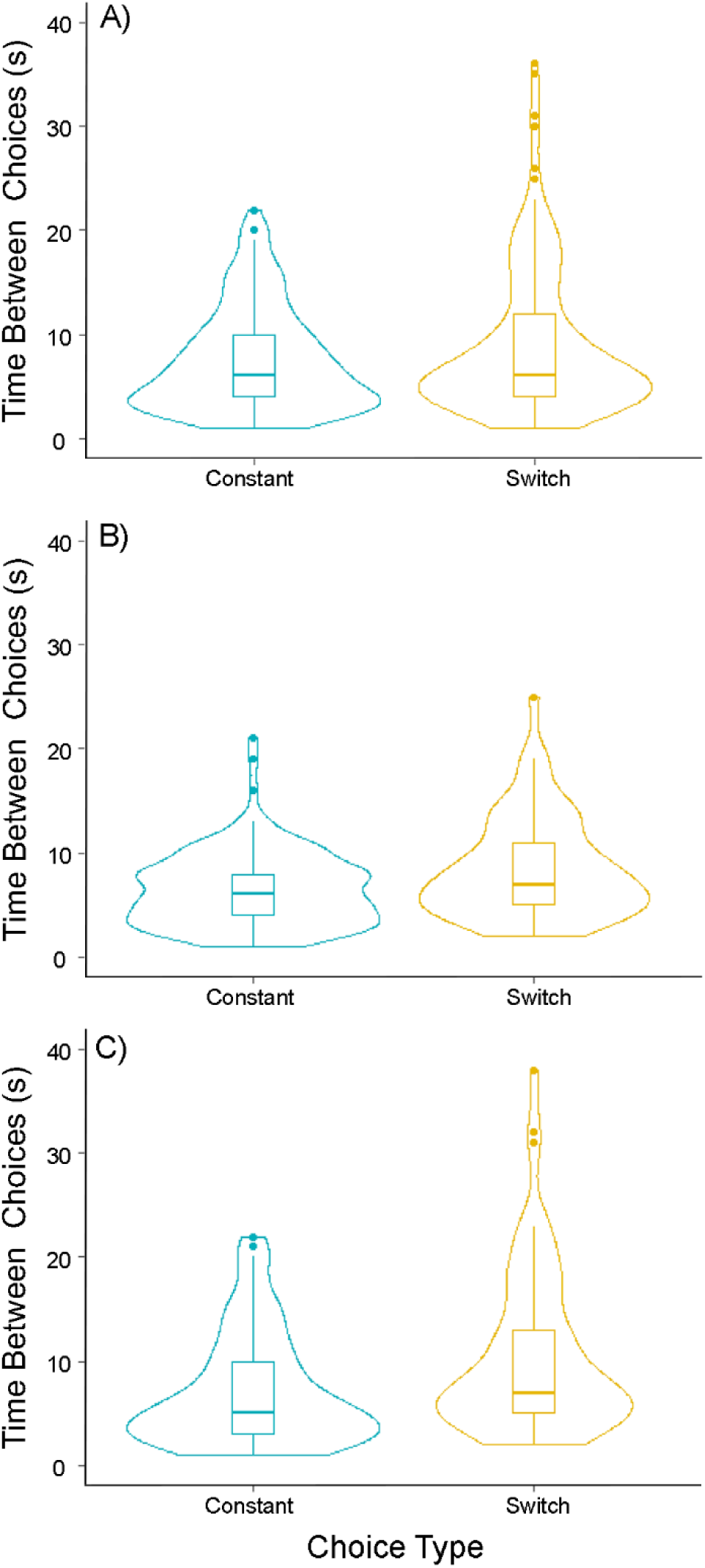
Time taken to make constant choices and switches in A) Experiment 1, B) Experiment 2, and C) Experiment 3. Other details as for Figure 2.

### Experiment 2: How does reward value influence bee visual search?

The average time taken for the first and second training bouts on this experiment was 1033.8 (± 439.8) seconds and 958.7 (± 493.4) seconds respectively.

The average proportion of high-reward targets chosen by bees was 0.69 (± 0.18 S.D.), while the average proportion of low reward targets chosen was 0.28 (± 0.17 S.D.). The best model for the proportion of choices included an interaction between the reward value and the order of the training. Higher reward value (50% Sucrose) led to significantly greater proportion of choices compared to both low reward value (30% Sucrose) flowers (GLM, effect size estimate: −1.12, P = 2.15*10^−5^, Fig 2B) and distractors (GLM, effect size estimate: −3.65, P = 1.18*10^−13^). Thus, bees chose high-reward targets more often than low-reward targets. The average proportion of choices made to distractors was 0.02 (± 0.04 S.D.), demonstrating that the bees were capable of simultaneously choosing between two targets even in the presence of distractors.

Bees that were first trained on high-reward targets chose these targets significantly less than if they were first trained on low-reward targets (GLMM, effect size estimate: 0.72, P = 0.0088). There was also a significant interaction effect between training order and reward value (GLMM, effect size estimate: −1.38, P 0.00041). Bees were thus more likely to choose high-reward targets if they had been trained on them in the bout immediately preceding the test (i.e. trained on the low-reward targets first, yellow/right vs blue/left plots in Fig 2B). The training times between the end of the first bout and the start of the test were, however, not significantly different when the first training bout had targets of high or low reward value (Wilcoxon rank sum test, W= 29, P=0.57). The interaction effect between training order and reward value is thus not due to difference in training times.

The average sequence index of the bees was 0.69 (± 0.20 S.D.) and this was significantly different from 0.5 (Wilcoxon rank sum test, W = 187.5, P = 0.0008, Fig 3). This indicates that in this experiment bees were more likely to have constant choices than switches. The time taken between choices was also significantly different between constant choices and switches chosen (Wilcoxon rank sum test, W = 2661.5, P = 0.01, Fig 4B). The mean time taken for constant choices was 6.49 (± 3.63 S.D.) seconds compared to a mean of 8.47 (± 4.88 S.D.) seconds for switches.

### Experiment 3 How does bee visual search combine reward value and physical saliency?

The average time taken for the first and second training bouts on this experiment was 1884.5 (± 993) seconds and 1681.1 (± 815.3) seconds respectively.

The average proportion of high-reward, low-saliency targets chosen by bees was 0.56 (± 0.27 S.D.), while the average proportion of low-reward, high-saliency targets chosen was 0.34 (± 0.26 S.D.). There was no significant main effect of reward value on the proportion of high and low reward targets chosen (GLM, effect size estimate: 0.32, P = 0.23, Fig 2C) but a significantly higher proportion of high reward targets were chosen compared to distractors (GLMM, effect size estimate: −2.42, P = 2.55 * 10^−8^). Thus, bees chose high-reward targets as often as low-reward targets, despite their lower saliency. The average proportion of choices made to distractors was low at 0.10 (± 0.12 S.D.), demonstrating that the bees were capable of simultaneously choosing between two targets even in the presence of distractors.

The order in which bees were trained on the high-reward and low-reward targets had a significant main effect (GLM, effect size estimate: 1.1654, P = 2.01 * 10^−5^). There was also a significant interaction effect between reward value and the order of the training (GLM, effect size estimate: −2.8688, P = 3.39 * 10^−12^). Bees were thus more likely to choose high-reward targets if they were the targets in the second training session (immediately prior to the test) rather than in the first training session.

The training times between the end of the first bout and the start of the test were however not significantly different when the first training bout had targets of high or low reward value (Wilcoxon rank sum test, W= 31, P=0.78). The interaction effect between training order and reward value is thus not due to difference in training times.

The average sequence index of the bees was 0.65 (± 0.25 S.D.) and this was significantly different from 0.5 (Wilcoxon rank sum test, W = 192, P = 0.0084, Fig 3). This indicates that in this experiment, bees were more likely to have constant choices than switches. The duration between choosing one flower and the next was also significantly different between constant choices and switches (Wilcoxon rank sum test, W = 3184, P = 0.00053, Fig 4C). The mean time taken for constant choices was 7.14 (± 5.36 S.D.) seconds compared to a mean of 10.51 (± 7.78 S.D.) seconds for switches.

The mean search time spent before choosing a high-reward flower was 7.07 (± 5.15 S.D.) seconds while the mean search time spent before choosing a low-reward flower was 9.51 (± 7.33 S.D) seconds, and these values were significantly different (GLM, Estimate = −0.009, P=0.009). Thus, the bees were quicker to choose high-reward targets compared to low reward targets. The model that best explained the proportion of time bees spent in different zones in the arena included flower type and the order in which bees were trained on high or low reward flowers as factors. Bees spent a significantly greater proportion of time inspecting higher rewarding flowers than lower rewarding flowers with greater physical saliency (GLMM, effect size estimate = −0.63, P < 2 * 10^−16^ Fig 5A) and distractors (GLMM, effect size estimate = −2.14, P < 2 * 10^−16^). There was also a significant main effect of the order in which bees were trained on high or low reward flowers (GLMM, effect size estimate = 0.84, P < 2 * 10^−16^) as well as an interaction effect between flower type and the order of training (GLMM, effect size estimate = −2.28, P < 2 * 10^−16^). Thus, when bees were trained on the high reward flowers first and the low-reward flowers later, they were equally likely to spend time around high-reward, low saliency flowers and low-reward high-saliency flowers. However, when trained on the low-reward flowers first and the high-reward flowers later, they spend a greater time around high-reward low saliency flowers compared to low-reward high-saliency flowers.

**Fig. 5.**
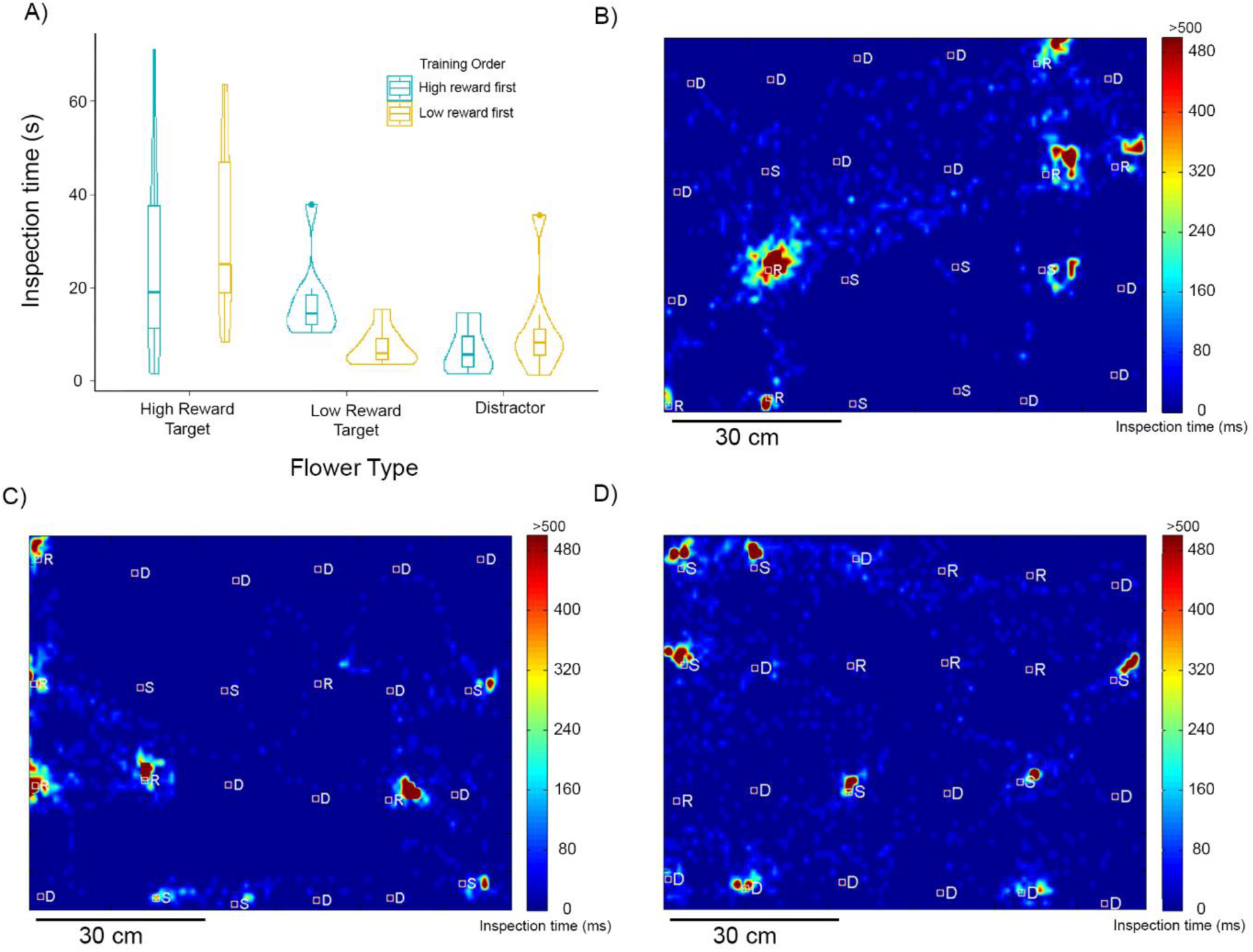
Inspection times around different flower types. Details for A) as in Figure 2. B-D) Example visual search maps for three bees depicted as a top view of the flight arena with targets and distractors. Colours depict the inspection times up to a maximum of 500 ms (only 5% of all times were greater than this limit). Squares depict flower positions. R = High-reward, low saliency targets; S = Low-reward, high-saliency targets, D = Distractors. B and C depict examples were bees spent more time around high-reward targets, D depicts an example where the bee spent more time inspecting low-reward targets.

## Discussion

Bees trained on multiple targets can choose the targets in the presence of distractors, without staying flower constant (Nityananda and Pattrick 2013). When targets are matched in both saliency and reward, bees are equally likely to choose either rewarding target, and switch between them often. Our results here demonstrate how bees prioritize learnt rewarding targets when they differ in physical saliency, reward value or both. We find that differences in saliency and reward value do not hamper the visual search task and bees are still able to choose at least two target types and ignore distractors. Both saliency and reward influence the proportion of targets chosen – with more salient and more rewarding targets chosen in higher proportions. The order in which bees encounter the targets during training matters when the targets differ in reward value and bees show a recency effect (Ebbinghaus 1885). This is particularly evident when the targets differ in both saliency and reward value. While bees in this condition seem to choose high reward low-saliency targets at an equal proportion as low-reward high-saliency targets, a slightly different pattern is seen when training order is accounted for. Low-reward, high-saliency targets are more likely to be chosen if they are encountered in the most recent training bout rather than the earlier training bout. This effect is less pronounced for the high-reward, low saliency targets. Our results also show that high-reward targets lead to greater flower constancy, shorter times for constant choices and more time spent attending to these targets.

Studies have long shown that bees can differentiate between coloured targets that differ in reward value (Lubbock 1881; Turner 1910; von Frisch 1914; Benard et al. 2006; Avarguès-Weber and Giurfa 2014). Most studies, however, have typically used appetitive training paradigms where bees are trained to distinguish targets with a reward from distractors without a reward (Avarguès-Weber and Giurfa 2014). More recently, studies have focussed on aversive training paradigms where bees distinguish between targets that are rewarding and distractors that contain an aversive solution like quinine (Dyer and Chittka 2004b; Giurfa 2004; Avarguès-Weber and Giurfa 2014). These two approaches have different effects with aversive conditioning leading to more fine-grained colour discrimination (Dyer and Chittka 2004b; Giurfa 2004). Studies that use two stimuli that are both rewarding but differ in reward value, as in this study, are fewer but they clearly demonstrate that bees can learn to differentiate colours even in this paradigm (Baude et al. 2011; Riveros and Gronenberg 2012; Avarguès-Weber et al. 2018). In one study using harnessed, rather than free-flying bees, the reward differential was provided by either providing the same concentration of sucrose solution to both the antenna and the proboscis (high reward condition) or to only the antenna (low reward condition). This differential was sufficient for bees to distinguish the colours associated with higher reward from those associated with lower rewards (Riveros and Gronenberg 2012). Our results from experiment 1 demonstrates that flowers that have a higher sucrose concentration are preferred by freely flying bees and bias their visual attention. The results from experiment 3 further show that flowers previously associated with high reward are still chosen half an hour after the training, even when they have lower saliency than low reward flowers.

The influence of physical saliency or colour contrast on bee visual search is less well studied than the influence of reward value (but see (Spaethe et al. 2001)). However, some studies have looked at this in the context of the innate preferences of bees (Lunau 1990; Giurfa et al. 1995; Lunau et al. 1996). These preferences are typically biased towards the UV-blue spectral range but do not seem to reflect the colour or green contrast difference from the background (Giurfa et al. 1995). Flower colours that have high spectral purity against background with low spectral purity do however attract the strongest innate behavioural responses from bumblebees (Lunau 1990). In addition, while bees can be trained to overcome their initial biases, their preferences can remain influenced by the effect of innate preferences (Gumbert 2000). In our experiment 3 we used a blue target as a low reward target to see if the high reward value of the other target could overcome biases towards this target. We found this to occur if the bees were trained on the blue targets further in time from the test. Higher reward also biased visual attention away from the high saliency blue targets as indicated by the time spent by the bees around different types of flowers.

Our results also show that the search history of the bees is important to consider. Bees might often specialize on the first colour they find to be rewarding – regardless of saliency. This would prevent them from learning multiple targets as in our study. In fact, other studies have found persistent flower constancy when bees are not allowed to learn both targets independently (Wells and Wells 1983; Hill et al. 1997). In nature, multiple targets might possibly be learnt when floral communities are more diverse or have higher densities of flowers (Heinrich 1979; Gegear and Thomson 2004; Baude et al. 2011). Our results and those of previous papers show that bees can switch between flowers and do not always stay flower constant. Thus, flower constancy does not stem from a cognitive limitation as has been suggested before (Waser 1986; Raine and Chittka 2007). Our results further point towards the importance of reward value for constancy. Bees show greater flower constancy when the targets differed in reward value. In these cases, they also showed shorter times when making constant choices rather than switching between colours. Bumblebees have been shown to fly shorter distances after visiting rewarding flowers compared to non-rewarding flowers (Dukas and Real 1993). Our results show that the experience of different reward values could also influence their foraging behaviour. Bees appear more likely to switch between flowers that have equal reward value but stay constant to highly rewarding flowers. Flower constancy is also affected by the density of conspecifics (Baude et al. 2011) so including this along with reward value and perhaps floral diversity would make for a fuller picture of the ecology of flower constancy.

Reward value also appears to influence the visual attention of the bees in addition to constancy and choice latencies. Bees spent longer inspecting high-reward flowers compared to low-reward flowers of greater saliency and were quicker to choose them. This resembles results from the human visual search literature, especially experiments demonstrating that the reward value associated with a stimulus can influence reaction times even if the stimulus is not task-relevant or salient (Anderson et al. 2011a, b). In our experiments we cannot assign task goals to the bees. However, the training order serves as a proxy for this. Half the bees in experiment 3 were initially trained on the high reward target and then on the low reward target. When faced with the test, the most recent training could arguably be considered the relevant task, making the previous high-reward targets irrelevant stimuli. Nonetheless bees still chose and attended to these targets – paralleling results in human experiments. We might potentially see different results when the reward values are lower, or the contrast of the high reward target is reduced even further. When high-reward targets have very low detectability, low-reward targets with high physical saliency could have lower search times. In these cases, bees might then change their preference to low-reward targets rather than high-reward ones, especially if the rewards are not very different. It has been argued that reward-based attentional capture in humans arises from Pavlovian mechanisms, where the level of reward determines the effectiveness of attentional capture (Bucker and Theeuwes 2017; Mine and Saiki 2018). Since several animals, including bees, are well known for Pavlovian learning, we should therefore expect this form of attention to be widespread in several animals. Our results suggest this might be true in bees and more focussed experiments showing that the mechanisms of attentional capture are shared in bees and humans would be an exciting area for future research.

## Acknowledgements

VN is funded by a Biotechnology and Biological Sciences Research Council David Phillips Fellowship BB/S009760/1 and this work was partly carried out with support from a Marie Curie Incoming International Fellowship (PIIF-GA-2009-253593). We thank Sara Khan, Muhammad Sayyidul Hasan and Piranavan Selvaratnam for help with running the experiments.

