## Supplementary Table 1 for "Reward Value Is More Important Than Physical Saliency During Bumblebee Visual Search For Multiple Rewarding Targets"

**Electronic Supplementary Material**

Corresponding author: Vivek Nityananda

**Table S1:** Distance in hexagonal units between the different colours used in the experiments, as well as betweeen the colours and the background used in the experiments.

| | **Blue** | **Cream** | **Red** | **Yellow** | **Fuchsia** | **Green Background** |
|---|---|---|---|---|---|---|
| **Blue** | 0.00 | 0.13 | 0.42 | 0.41 | 0.31 | 0.45 |
| **Cream** | 0.13 | 0.00 | 0.43 | 0.38 | 0.35 | 0.47 |
| **Red** | 0.42 | 0.43 | 0.00 | 0.14 | 0.15 | 0.04 |
| **Yellow** | 0.41 | 0.38 | 0.14 | 0.00 | 0.25 | 0.18 |
| **Fuchsia** | 0.31 | 0.35 | 0.15 | 0.25 | 0.00 | 0.16 |
